# Spatially distributed growth factor environments for evaluating hydrogel-based CD34^+^ cell culture conditions

**DOI:** 10.64898/2026.09.15.751891

**Authors:** Ashley R. Murphy, Rose Ann Franco, Phillip Johnson, Justin J. Cooper-White, Mark C. Allenby

## Abstract

Suspension culture is the gold standard for *ex vivo* hematopoietic stem cell (HSC) expansion; however, high media requirement results in expensive cell therapy products. Hydrogel-based HSC culture represents a more realistic *in vitro* representation of human *in vivo* HSC microenvironments and may support a more efficient use of media during *ex vivo* HSC expansion. However, media conditions used for hydrogel-supported cell culture often simply mimic that which are currently used in suspension culture approaches, potentially resulting in an oversupply of cell growth proteins. In this work, we present a microfluidic culture system capable of supporting hydrogel-based cell culture and generating spatial distributions of independent growth factor combinations. Using this device, we image, in real-time, the behaviour of human umbilical cord blood-derived CD34^+^ cells, within fibrin-based hydrogels, in response to spatial gradients of the human recombinant growth factors stem cell factor, thrombopoietin, angiopoietin-2 and insulin-like growth factor-II. Using live microscopy and image-based single-cell segmentation and quantification, local changes in cell density over time in relation to spatial concentration of factors were identified. This novel microfluidic device has the potential to screen combinations of growth factors required for hydrogel-supported CD34^+^ cell expansion and inform culture media formulations of future hydrogel-based cell manufacturing processes.

**Table of Contents Figure**

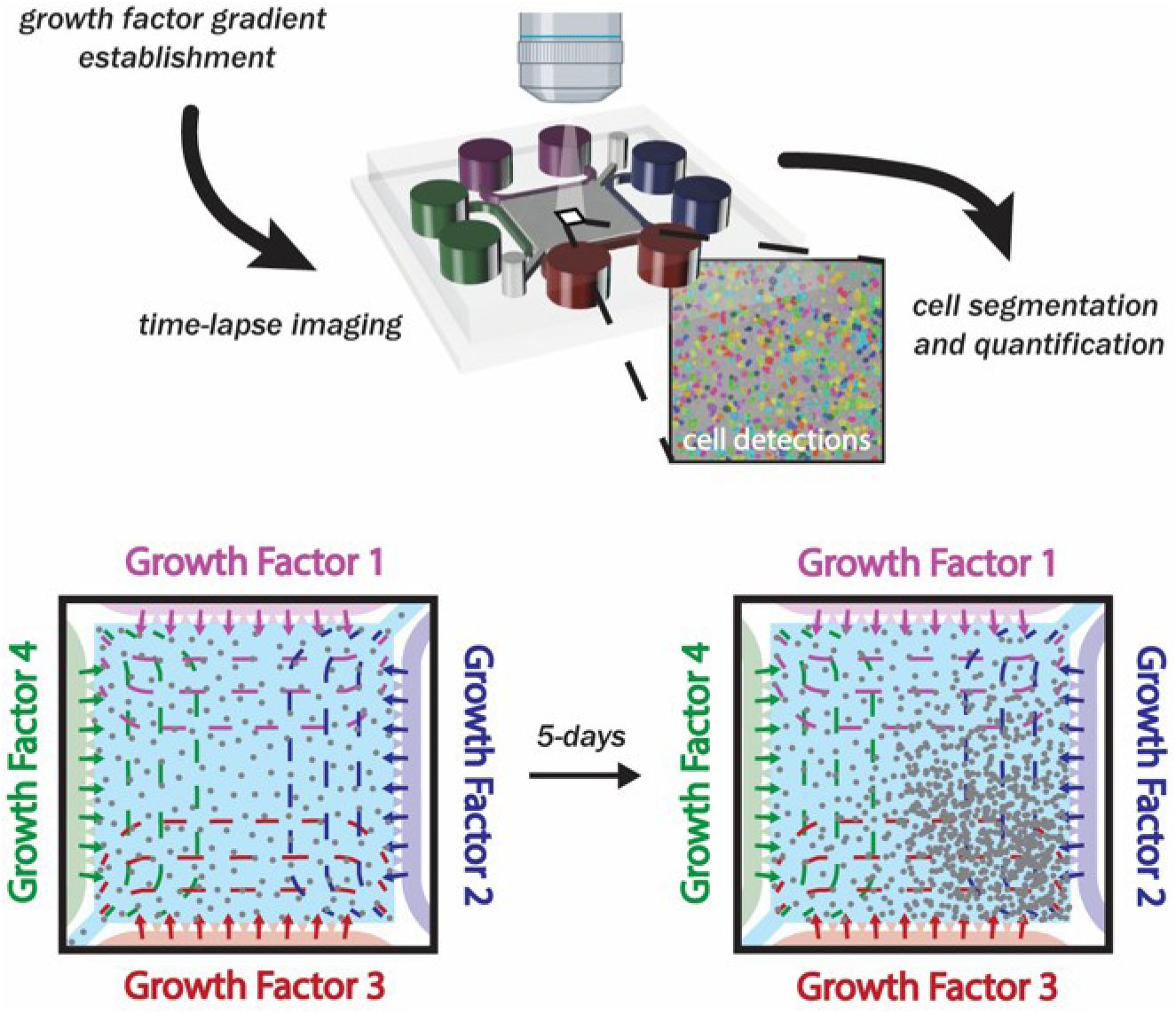

**Table of Contents Text**

Microfluidic devices supported the establishment of independent growth factor gradient distributions across a fibrin hydrogel hematopoietic stem cell culture. Time-lapse live cell imaging, cellular segmentation and quantitative analysis demonstrated spatially preferential hematopoietic stem cell expansion. This technology demonstrates the potential to evaluate supplemented growth factor type and concentration in supporting hydrogel-based CD34^+^ cell culture conditions.

## 1. Introduction

The capacity of hematopoietic stem cells (HSCs) to reconstitute the entire human immune system following transplantation has been known and leveraged for decades as a potential curative treatment option for patients diagnosed with a range of blood disorders including leukemia, metabolic storage disorders, immunodeficiencies and bone marrow failure syndromes. However, in recent years their potential to serve more broadly in the context of next generation cellular therapies has emerged. While bone marrow and mobilized peripheral blood provide a rich source of HSC, clinical procedures to obtain this biological starting material are invasive and not without real risk to the donor. In contrast, allogeneic umbilical cord blood (UCB)-derived HSCs overcome these risks and represent a promising alternative cell source. Consequently, 13 of the 46 cell and gene therapies currently approved by the US Food and Drug Administration include UCB-derived HSCs.^[1]^

Since its first use in the HSC transplantation (HSCT) setting in the 1980’s UCB has faced challenges with its use.^[2]^ Most notable is the limited HSC cell dose/kg patient body weight contained in any single donation, which can range from 2.3–18.3 × 10^6^ CD34^+^ cells per donation.^[3]^ Current HSCT practice uses ≥1.5 × 10^5^ CD34^+^ cells per patient kg.^[4]^ A 2019 study of 126,341 cord blood units contained within the US public cord blood bank cryopreserved inventory reported only 4% had sufficient total nucleated cell and CD34^+^ cell doses for a single-graft transplant of a patient of 70kg.^[5]^

In an attempt to overcome limited CD34^+^ cell numbers, researchers have looked towards ‘pooling’ donors, which increases pathogenic risk to the patient and likelihood of immune rejection, or expanded *ex vivo* at great financial expense and risk of lessened potency. *Ex vivo* cell expansion typically involves harvesting the UCB-derived mononuclear cell fraction, isolating the CD34^+^ cells and culturing them suspended in a cocktail of growth factors and small molecules supportive of cell maintenance and proliferation.^[6,7]^ As of 2025, there are 26 ongoing or concluded clinical trials exploring the safety and efficacy of *ex vivo* expanded CD34^+^ cells and currently, OMISIRGE® (omidubicel-onlv) by Gamida Cell Ltd remains the first and only market approved *ex vivo* expanded CD34^+^ cell-containing therapy.^[8,9]^

Typical e*x vivo* HSC expansion protocols adopt static or stirred suspended culture methods with densities as low as 10^4^ cells mL^-1^ in nutrient rich media with concentrations of clinical grade recombinant growth factors as high as 300 ng mL^-1^ for culture periods as long as 21 days, culminating in exorbitant media costs to expand one patient dose to >10^8^ HSCs.^[10]^ With the recent surge in approved UCB-derived HSC therapies, advances in *ex vivo* culture approaches to address current manufacturing limitations are required to rapidly suffice growing market demands.

We look to human physiology for inspiration in terms of design parameters and methodologies for expanding HSCs *ex vivo*. The fetal liver and adult bone marrow are the primary physiological niches which support expansion and maintenance of HSCs.^[11,12]^ Both provide an extracellular matrix which supports cellular organisation, sequesters and delivers bioactive molecules, facilitates cell-cell communication, provides physical forces that regulate HSC behaviour, maintains quiescence and encourages proliferation.^[13]^ Hydrogel matrices studied for the encapsulation of HSCs range from completely natural to fully synthetic and include collagen I, Matrigel®, fibrin, alginate, gelatin methacrylol, and polyethylene glycol diacrylate.^[14]^ Fibrin hydrogels have been shown to be particularly effective in maintaining and expanding CD34^+^ cells *ex vivo* and additionally have the future potential to be derived from UCB towards a fully patient-matched culture system.^[15–17]^

The presence of natural or synthetic hydrogels as extracellular matrix mimetics is known to alter HSC behaviour and function.^[14]^ This may be attributed to the direct application of culture parameters (e.g., media composition, volume, growth factor concentration and type) used in traditional liquid suspension cultures to this more physiologically representative culture method. Optimisation of hydrogel-based culture parameters may not only afford the opportunity to better understand and potentially resolve the challenges in maintaining post-expansion HSC function, but it also represents an opportunity to realise significant savings in laboratory resources and reduce the current high cost of manufacturing and supply.

To address this, we have developed a microfluidic culture system designed to generate a spectrum of concentrations of up to four independent growth factors across CD34^+^ hematopoietic stem and progenitor cells (HSPCs) encapsulated in fibrin hydrogel. The device configuration allows for real-time imaging of single-cell locations and morphologies to evaluate and optimise combinations of desirable growth factors resulting in maximal cell expansion. Computational modelling of growth factor transport-decay and quantitative image analysis of cultures allows cell expansion and location to be assessed against theoretical local growth factor concentrations, permitting elucidation of optimal growth factor combinations and concentration for future use in larger, scaled-out culture.

As a proof of concept, we identified four soluble factors to study: two established cytokines common to the bone marrow *and* fetal liver niches, stem cell factor (SCF) and thrombopoietin (TPO); and two thought to be *exclusive* to the fetal liver niche, angiopoietin-like protein 2 (ANG-2) and insulin-like growth factor II (IGF-II). SCF and TPO are well established factors which support and maintain HSC in *ex vivo* suspension culture and are present in most commercial HSC expansion media formulations, whereas both ANG-2 and IGF-II have limited literature validation for their influence on *ex vivo* HSC expansion.^[18]^ ANG-2 from freshly isolated fetal liver stromal cells has been identified to expand HSCs.^[19]^ Additionally, fetal liver stromal cells express the cytokine IGF-II, along with SCF and TPO, and have been shown to maintain the proliferative capacity of embryonic day 15.5 fetal liver HSCs for up to 4-days in co-culture.^[20]^ We herein demonstrate the capability of this device and the associated methodology to evaluate the influence of these growth factor combinations for supporting UCB-HSPC expansion within fibrin hydrogels.

## 2. Results and Discussion

### 2.1. Culture setup and optimization

When pipetted into the loading port of the microfluidic culture device, bovine fibrin hydrogel precursor solution-cell suspensions of 10 µL slowly filled the 7.64 × 7.64 × 0.05 mm culture compartment within approximately 5 seconds (figure 1. A–B, figure S1). Ejecting the pipette tip immediately after the solution entirely filled the culture compartment prevented the hydrogel precursor solution from flowing through the trapezoidal pillars and breaching confinement. The tip could then be carefully removed using a twisting motion while pulling upwards as not to disturb the culture. Trapezoidal pillars of dimensions 80 µm (short axis), 270 µm (long axis) and 200 µm height with a minimum spacing of 100 µm (figure S1) were found to adequately contain fibrin hydrogel precursor-cell mixtures and maintain containment post-gelation and throughout culture up to 5-days. Due to the low 50 µm height of the hydrogel compartment and high projected area (0.58 cm^2^), it was important to not apply any undue force to the surface of the device as this would force hydrogel out of the culture compartment and into the media channels. The two culture compartment corners adjacent to the hydrogel inlet and outlets often did not consistently fill completely (figure S2). While this was assumed to have no impact on culture experiments, future studies could remove these corners with a flat edge to support complete filling.

**Figure 1.**
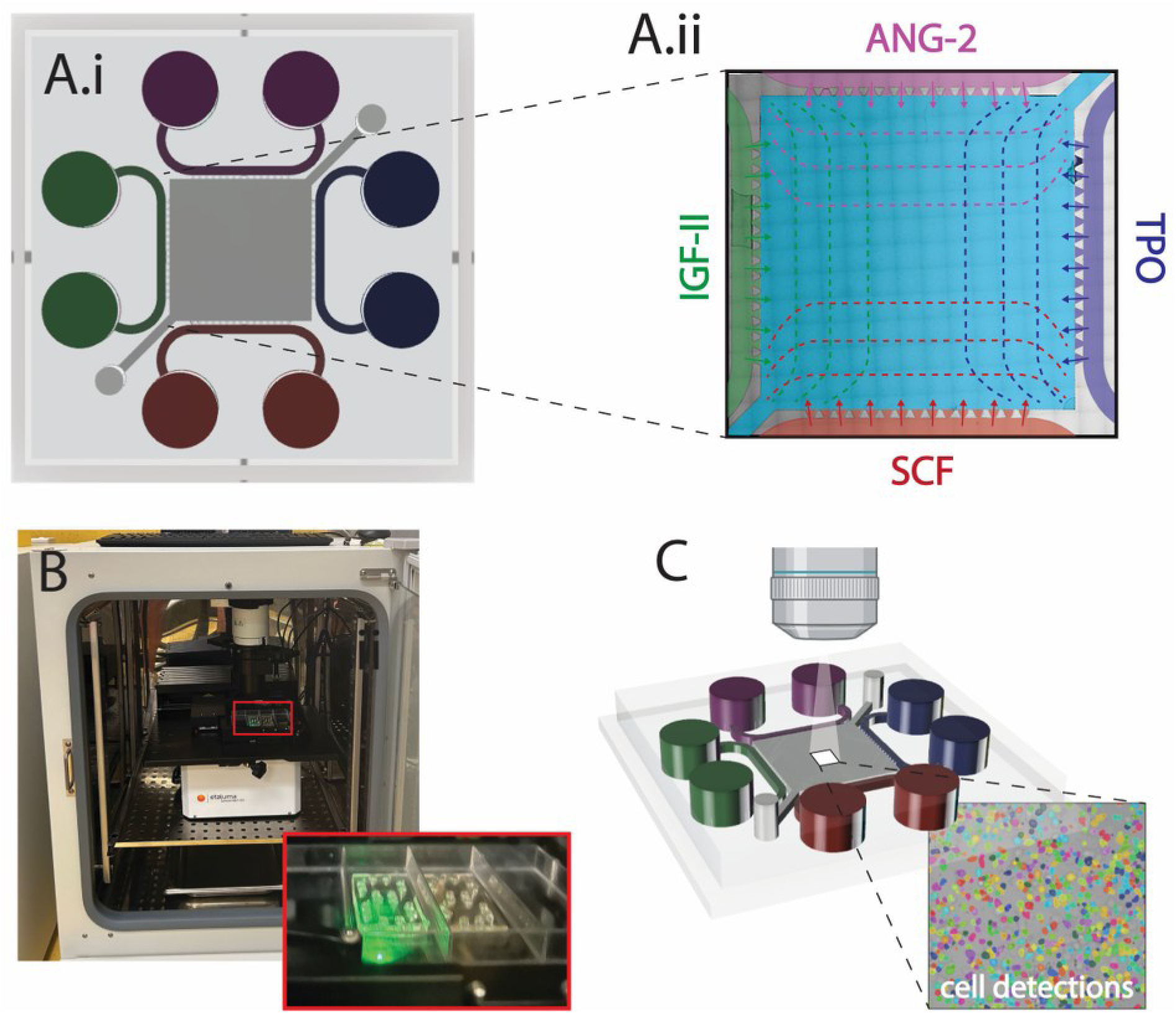
Four-channel microfluidic culture devices support the establishment of growth factor gradients across a square culture compartment and are amenable to live-cell imaging and cellular segmentation. **A.** A 7.64 × 7.64 mm culture compartment confines a hydrogel cell suspension (grey) using trapezoidal micropillars (*i*). Along the four edges of the culture compartment, growth factor solutions (purple, blue, red, green) are provided and diffuse across the hydrogel culture (*ii*). **B.** The devices are placed in a rectangular well plate and are subjected to brightfield time-lapse imaging using an in-incubator motorised stage microscope. **C.** Image data sets are subject to cell segmentation using the cellpose algorithm. ANG-2: angiopoietin-2, SCF: stem cell factor, TPO: thrombopoietin, IGF-II: insulin-like growth factor-II.

A fibrin concentration of 1 mg ml^-1^ visibly maintained hydrogel integrity throughout the 5-day culture and resulted in maintenance of a CD34^+^-like cell morphology throughout the first 24-hours post-seeding (video S1). A fibrin hydrogel concentration of 0.5 mg mL^-1^ was observed to visibly degrade throughout culture via the formation of liquid currents throughout the hydrogel (data not shown). It is suspected that the degree of crosslinking at 0.5 mg mL^-1^ fibrin is insufficient to withstand degradation resulting from CD34^+^ cells and their progeny, despite the presence of the anti-fibrinolytic enzyme, aprotinin. A fibrin hydrogel concentration of 3 mg mL^-1^ resulted in a sudden change in cell morphology within 24-hours, evidenced by a reduction in cell diameter and roughening of cell borders (data not shown). It is expected the mechanical properties of the 3 mg mL^-1^ fibrin hydrogel was too stiff to maintain CD34^+^ cell viability, as has been demonstrated in previous studies investigating the influence of extracellular matrix mechanical properties on CD34^+^ cell behaviour.^[21,22]^ Inclusion of 0.15 U ml^-1^ aprotinin in the hydrogel precursor mixture and in the liquid media was necessary to maintain hydrogel integrity and prevent degradation thought 5-days culture, as was the case when omitted from liquid culture media (data not shown).

An in-incubator fluorescence/brightfield microscope (figure 1.B) was found capable of imaging the entire culture compartment of three microfluidic culture devices in parallel. A 20× objective, in combination with the 50 µm culture chamber height sufficiently captured cellular detail necessary for image segmentation, whilst minimizing out-of-focus objects (figure 1.C, figure S2). An imaging grid of 15 × 15 tiles (1600 × 1600 pixels per tile, 0.419 µm per pixel) with 10% overlap was found to be appropriate for reconstructing an image of the entire culture compartment with visibly distinguishable cell borders (figure S2–3). Each independent complete culture compartment (15 × 15 tiles) was imaged on average every 8.66 minutes (1.30 seconds per image including stage movement). Two complete media changes were conducted during the 5-day cultures as close as practicable to 24- and 72-hours post seeding. The time duration to conduct media changes on three independent cultures and reset all 3 × 15 × 15 image coordinates was on average 3.34 ± 0.25 hours. When replacing the culture plate, the device culture compartment would often fall out of the microscope field of view, therefore the 15 × 15 regions of interest were reset after each media change. A dedicated device holder which returns the devices exactly back to the same imaging position post-removal could circumvent this issue and prevent downtime in imaging during future experimentation.

### 2.2. Cell expansion quantification

UCB-derived CD34^+^ cells were cultured using microfluidic devices in 1 mg ml^-1^ fibrin hydrogels at a seeding density of 10 × 10^6^ cell mL^-1^ in the presence of StemSpan SFEM II media (‘StemSpan’, in all channels), Stem Line II media with 100 ng ml^-1^ each of ANG-2, IGF-II, SCF and TPO (‘Stem Line II’, in all channels) and Stem Line II media with either 100 ng ml-1 ANG-2, IGF-II, SCF and TPO (‘gradient’, in each channel, respectively). The anatomical segmentation algorithm ‘cellpose’ was found to segment cells from completely stitched single plane brightfield images acquired starting from approximately 2-hours post seeding up to termination at approximately 5-days (average termination time 4.80 ± 0.23 days) with an accuracy of 93.86 ± 5.02% accuracy over 4,647 cells in 24 manually scored images at varied densities (figure S3). At a seeding density of 10^7^ cells mL^-1^, the 2.92 µL volume of the segmented culture compartment should theoretically contain 2.92 × 10^4^ cells. The first time point, which occurs approximately 2-hours post seeding, was found via cellpose detection to average 3.24 ± 0.74 × 10^4^ cells. Accounting for some cell expansion occurring during this culture period, the accuracy of the cellpose cell detection software seems to adequately detect correct total cell numbers within a reasonable margin.

All media conditions demonstrated an increase in global cell number and density at the conclusion of culture (figure 2.A–B, video S2–3). Discrete gradient cultures typically demonstrated a plateau in global cell density after approximately 48-hours of culture. Both StemSpan and Stem Line II conditions demonstrated an increase in cell density throughout the entire culture period and revealed a reduction in expansion rate after approximately 72-hours culture (figure 2.A–B). The StemSpan condition of donor 4 uniquely demonstrated a constant positive expansion rate throughout the entire length of culture (figure 2.A.iv).

**Figure 2.**
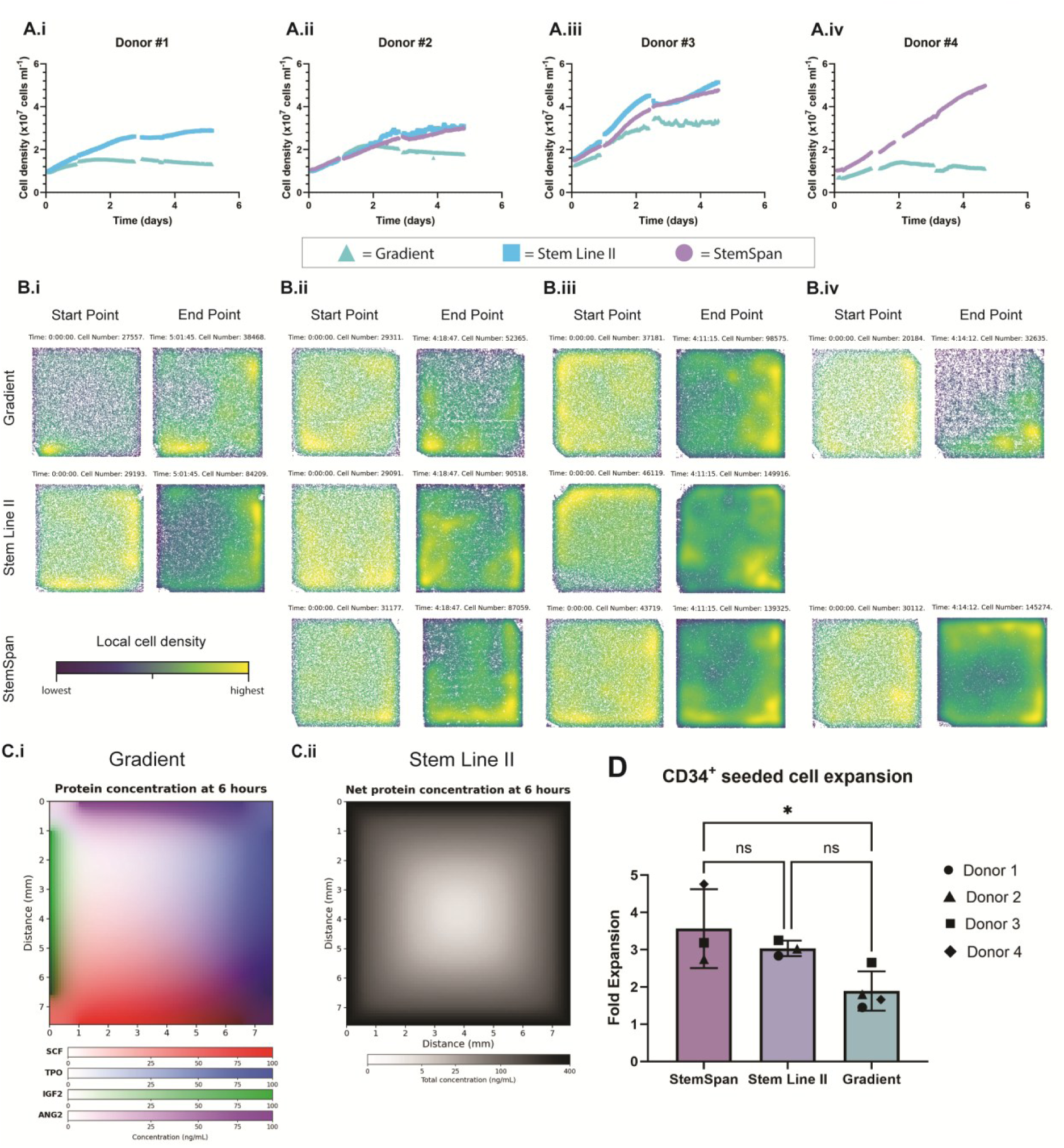
Fibrin hydrogel-based CD34^+^ cell cultures respond differentially to growth factor type and concentration when compared to control media mixtures. CD34^+^ cells seeded at a density of 10 × 10^6^ cell mL^-1^ in 1 mg ml^-1^ bovine fibrin hydrogels within a culture compartment volume of 2.92 µL and cultured up to 5-days. **A.** Cells detected via the cellpose software and plotted as global cell density over time for donors 1 to 4 (*i*–*iv*). Discontinuations in data represent culture media exchanges and resetting of the xyz-imaging coordinates. **B.** Global cell detection with qualitative representation of local cell density with dark blue representing the lowest local cell density in a particular culture image set and yellow representing highest local cell density, for donors 1 to 4 (*i*–*iv*). Images represent the first images (taken approximately 2-hours post-seeding) and the final image (taken approximately 120 hours post-seeding). **C.** Numerical simulations of independent (*i*) and combined (*ii*) growth factor distributions across the culture compartment after 6-hours. **D.** Fold expansion of initially seeded CD34^+^ cells calculated from the earliest image time point to the final image timepoint. CD34^+^: cluster of differentiation 34 positive, ANG-2: angiopoietin-2, SCF: stem cell factor, TPO: thrombopoietin, IGF-II: insulin-like growth factor-II.

### 2.3. Morphological evaluation

Upon seeding, cells were visibly observed to have a circular shape with consistently similar diameters and tightly defined borders (figure 6.C–G). Cells were observed to undergo cell division into two similarly sized cells, indicating mitotic activity (figure 6.C–D, video S1). Cell migration through the hydrogel was observed and when doing so, cells adopted an elliptical morphology before settling and returning to a spherical morphology (figure S3.E, video S1). Changes in morphology identified by a reduction in cell diameter were visibly observed (figure S.3.F–G, video S1). This change in morphology could be characteristic of cell differentiation or apoptosis. On occasions, cells were seen to eject intracellular material and subsequently reduce in diameter, potentially characteristic of erythrocyte lineage differentiation or apoptosis (figure S.3.F–G, video S1).

Cell diameter was quantified from cellpose supported segmentation results and subsequent image analysis (figure 3, table S1). At the commencement of imaging (approximately 2-hours post-seeding) initial cell diameters averaged 24.22 ± 16.99 µm, 23.04 ± 14.25 µm, 23.27 ± 14.20 µm, and 22.32 ± 12.84 µm which uniformly decreased during the 5-day culture into final cell diameters of 22.05 ± 15.96 µm, 17.46 ± 13.06 µm, 17.75 ± 12.57 µm, 19.56 ± 13.35 µm for donors 1, 2, 3, and 4, respectively. This decrease appeared to be driven by the gradual decline of a larger 22 µm diameter cell and the emergence of a smaller cell of approximately 14 µm for all donors except donor 1.

**Figure 3.**
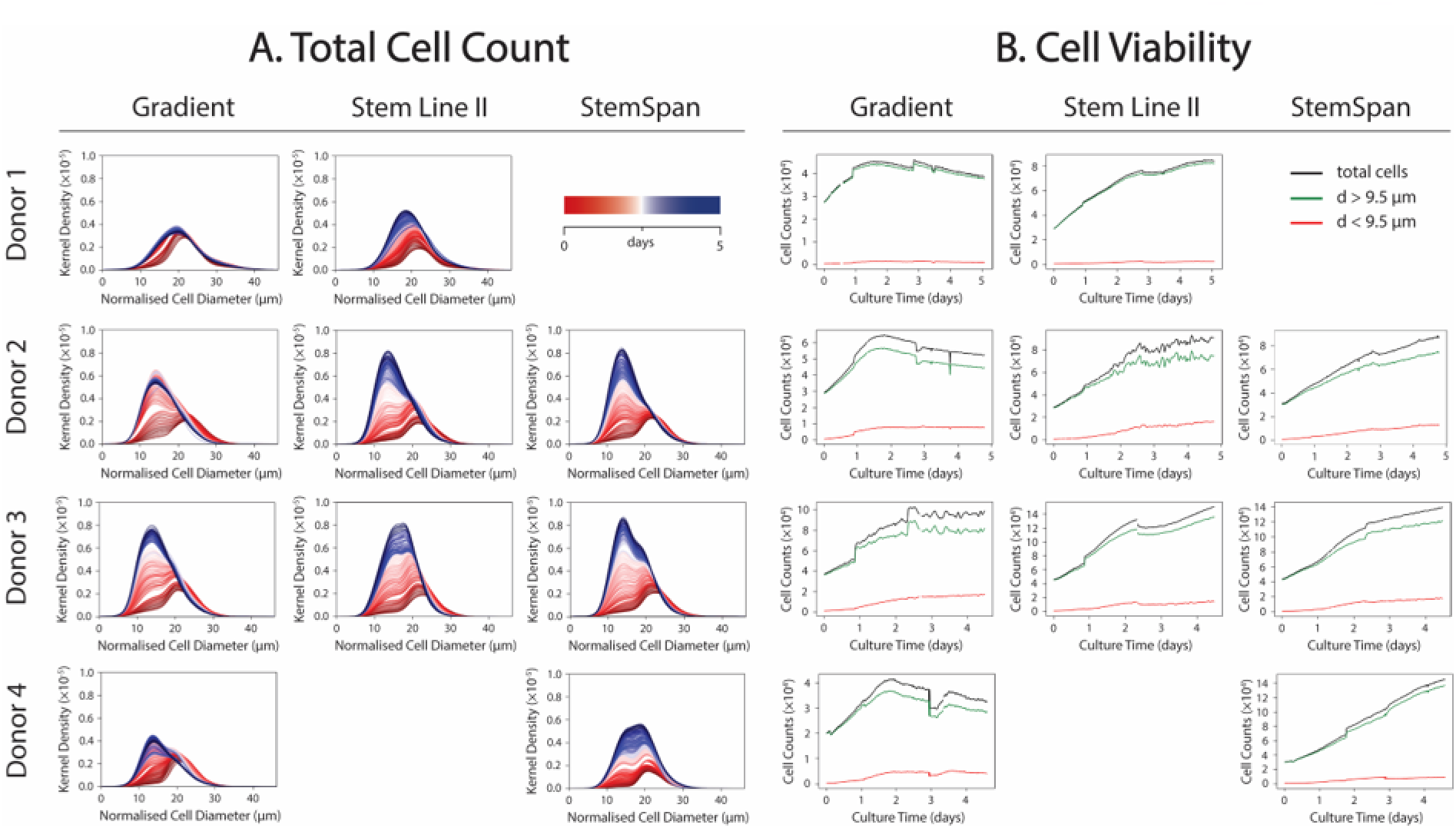
CD34^+^ cells demonstrate a change in cell diameter throughout the 5-day fibrin hydrogel-based culture. **A.** Kernel density distributions are plotted as a function of normalised cell diameter across the 5-days of culture (red t = 2 hours, blue t = 5 days) for the Gradient, Stem Line II and StemSpan conditions. **B.** Cell viability is estimated via a diameter threshold of 9.5 µm. Viable cells (d > 9.5 µm) are represented in green, non-viable cells (d < 9.5 µm) and total cells in black for the Gradient, Stem Line II and StemSpan conditions. CD34^+^: cluster of differentiation 34 positive.

During manual scoring of images to determine cellpose accuracy, it became apparent that later timepoints in less proliferative cultures contained even smaller objects that appeared to be fragmented non-viable cell membranes. This led us to apply the cellpose software to estimate culture viability by classifying smaller cells as putative ‘nonviable cells’. We set a nominal diameter of 9.5 µm as a viability threshold, in-line with our manual scoring and literature describing the size of apoptotic bodies, shrunken necrotic cells, or free-floating nuclei and debris. This arbitrary 9.5 µm threshold estimated culture viabilities of near-100% at the start of culture which were maintained near 98% for donor 1 and 4’s StemSpan configuration and declined to 90% for most other condition by the end of culture. Other similar ‘viability’ cell size thresholds could be user-selected in this platform.

It is important to note that determined HSPC sizes (22 µm at day 0) are nearly double those reported in literature.^[23]^ Therefore, it is likely cellpose consistently over-segmented cell areas. Therefore, should we have begun with a more realistic HSPC size of 11.5 µm, then smaller cells emerging towards the end of culture would average 7 µm (similar as reticulocytes or erythrocytes) and our 9.5 µm threshold for viable cell size more realistically represents 5 µm, smaller than all haematopoietic cell sizes.^[24]^

### 2.4. Response to growth factor gradients

Numerical modelling was used to predict the distribution of ANG-2, IGF-II, SCF or TPO across the 7.64 × 7.64 mm culture compartment (figure 2.C, figure S4). Simulations predicted these HSPC factors existed above 50% of their literature effective concentrations (EC50s) across 7% (IGF-II), 20% (ANG-2), 60% (SCF), and 100% (TPO) of the 7.64 mm square chamber. While a useful prediction tool, our simulations assume growth factors diffuse and decay similarly as found in liquid cell culture media and do not consider how CD34^+^ cell-laden fibrin hydrogels release, bind, or otherwise alter growth factor kinetics. The accuracy of quantitative computational predictions of growth factor transport could be improved with experimental measurements of growth factor reaction-diffusion within fibrin hydrogels. Nonetheless, the microfluidic chip culture designs should facilitate the formation of substantial growth factor gradients above and below bioactive concentrations, across the *x*- and *y*-axes of the culture compartment.

Cell density was assessed across the *x*- and *y*-axes of the culture devices and plotted over the entire length of culture (figure 4). Stem Line II and StemSpan conditions primarily demonstrated a symmetrical distribution of cell densities across both *x*- and *y*-axes. The Gradient conditions primarily demonstrated an asymmetrical distribution of cell densities across with *x*- and *y*-axes, with some potential bias towards the boundaries of the device containing the growth factors SCF and TPO, respectively.

**Figure 4.**
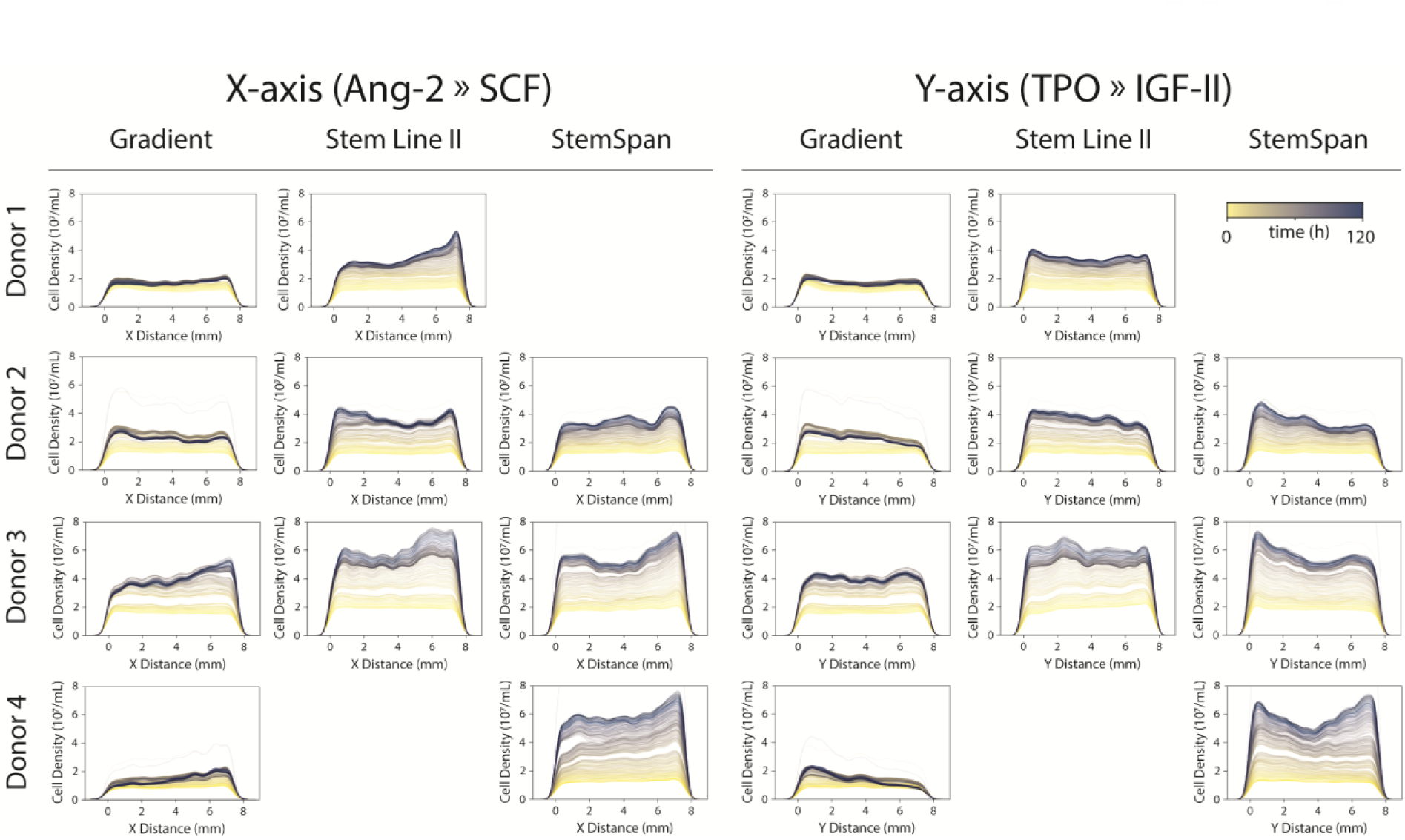
Global CD34^+^ cell expansion responds to discrete growth factor gradients. Cell density as a function of x-axis distance, with x = 0 representing the ANG-2 side of the culture compartment and x = 8 mm representing the SCF side. Cell density as a function of y-axis distance, with y = 0 representing the TPO side of the culture compartment and y = 8 mm representing the IGF-II side. Measurements taken across the entire 5-day culture period (yellow t = 2 hours, blue t = 5 days) CD34^+^: cluster of differentiation 34 positive, ANG-2: angiopoietin-like protein-2, SCF: stem cell factor, TPO: thrombopoietin, IGF-II: insulin-like growth factor-II.

### 2.5. Phenotypical characterisation

Cultures were fixed and immunocytochemically stained for the hematopoietic stem cell marker CD34, the mature erythroid lineage marker CD235a, the hypoxia responsive protein HIFa and the nuclear counterstain DAPI (figure 5, figure S5–S6). Despite significant staining time including agitation, the nuclear counterstain DAPI and the respective antibodies did not appear to diffuse sufficiently from the media channels entirely to the centre of the culture compartment (figure S5). To reduce the distance required for antibody diffusion, a future device should consider a removable top piece or film to allow staining solutions access from the top, and therefore only requiring a diffusion distance of 50 µm. Due to this limitation, high magnification images were therefore only collected at the four edges of the culture compartment (figure 5, figure S6).

**Figure 5.**
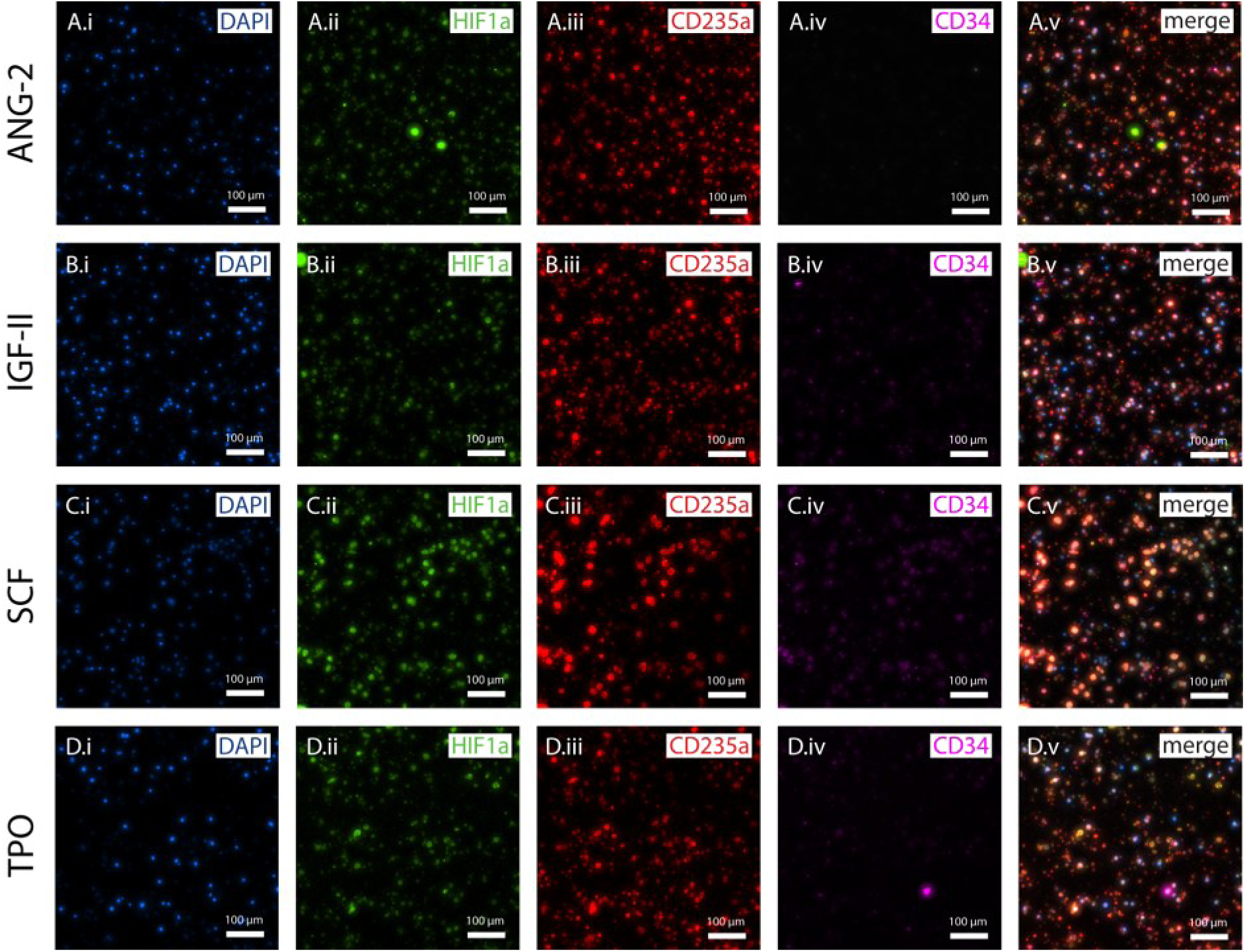
Fibrin hydrogel-based cells express hematopoietic lineage markers after 5-days culture in gradient conditions. Fibrin hydrogel-based cultures were fixed at day-5 post seeding and nuclear counterstained with (*i*) DAPI and immunocytochemically stained for the expression of the antigens (*ii*) HIF1a, (*iii*) CD235a and (*iv*) CD34. Images were captured directly adjacent to the (a) ANG-2, (b) IGF-II, (*iii*) SCF and (*iv*) TPO sides of the culture compartment. Isotype control stains can be found in figure S5 and S7. Scale bars = 100 µm.

Cells on all sides of the gradient microfluidic culture device strongly expressed the erythrocyte precursor marker, CD235a and weakly expressed the hematopoietic stem/progenitor marker, CD34 (figure 5.A.iii–iv, Biii–iv, Ciii–iv and D.iii–iv). Cells positive for the hypoxia responsive marker HIF1a were present on all sides of the culture device (figure 5.A.i, B.i, C.i, D.i). indicating some form of cellular stress occurring in the growth factor limited hydrogel environment. CD34^+^ cells initially seeded into the culture device suspended in fibrin hydrogel appear to lose their expression of the antigen, suggesting spontaneous differentiation in these limiting culture conditions and consistent with decreases in tracked cell diameter (figure 3.A).

### 2.6. Positional cell tracking

Cell-tracking is a common extension to cell culture platforms with integrated real-time imaging. While this hardware-software platform allows observations, in real-time, of the impact of growth factor gradients on high-cell-density hydrogel expansion culture cell number, density, shape, and/or phenotype, it does not elucidate whether these impacts were caused by cell proliferation, migration, or death. The ability to track individual live cell trajectories could determine whether zonal increases in cell density correspond to a particular growth factors ability to support of cell survival (by tracking cell death elsewhere), proliferation (new cells tracked in this zone), or migration (cells tracked to move from other zones, into this zone).^[25–27]^

This current platform currently struggles to integrate cell tracking, as scanning 15 × 15 tiles for each of two microchips (for donor 4) required 8 minutes, above the 1-to-5-minute single-cell tracking imaging rates found in other approaches.^[28]^ Furthermore, since the cells in this study are cultured in a 50 µm-high three-dimensional hydrogel; cells will likely migrate above or below the microscope focal plane. While future tracking approaches would benefit from a smaller length, width, and height culture chamber for more rapid and in-plane imaging of cells, as an exploration of this concept, live-tracking for donor 4 StemSpan condition was attempted (figure 6). For each of the 20,000–150,000 of cells detected within each of the 300–500 imaged timesteps, it was examined whether the next timestep presents a cell inside of a 150 µm Euclidean distance radius of that cell in the prior timestep. If multiple future cells were detected to be close to that past cell, the closest future cell was selected as the past cell’s next tracked position. If no future cells were detected in that threshold radius, the cell was considered to have died. If a new cell was detected and not related to a past cell, it was considered a newly proliferated cell.

**Figure 6.**
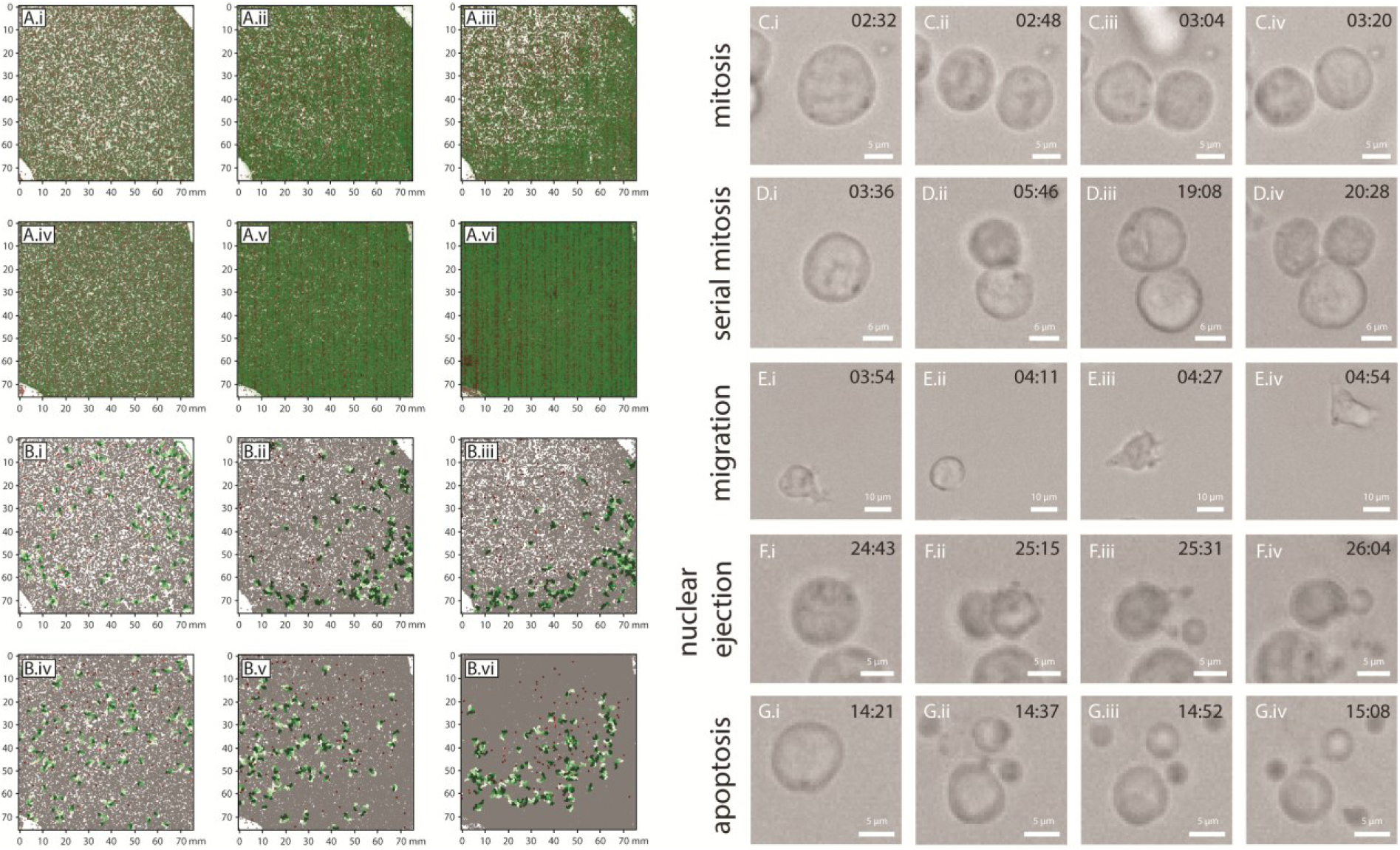
Single-cell-tracked proliferation and migration occurs along a SCF and TPO gradient. **A.** New cells tracked (proliferated; green) or lost (dead; red) for donor 4’s gradient (*i–iii*) and StemSpan (*iv–vi*) culture, from seeding to the first media change (*i*, *iv*), between the first and second media changes (*ii*, *v*), from the second media change to the culture conclusion (*iii*, *vi*). **B.** Of cells tracked for more than 100 time-steps, the top 100 moving (green) or stationary (red) cell tracks for donor 4’s gradient (i–iii) and StemSpan (*iv–vi*) culture, from seeding to the first media change (*i*, *iv*), between the first and second media changes (*ii*, *v*), from the second media change to the culture conclusion (*iii*, *vi*). All cells directly after media exchanges are pictured as grey in the background. Morphogenic events capturing mitosis (C), serial mitosis (D), cell migration (E), nuclear ejection-(F) and apoptotic-like events (G). Top left, time (hours:minutes post-seeding).

This cell-tracking approach identified that many new cells were generated via proliferation (green) close to the SCF and TPO growth factor gradients in donor 4’s gradient microchip (figure 6.A.i–iii) while proliferation in the StemSpan microchip occurred everywhere (figure 6.A.iv–vi). Also, this approach identified that, of cells tracked for more than 50 timesteps, cells near SCF and TPO growth factor gradients in donor 4’s gradient microchip were actively migrating after the first and second media changes (green tracks in figure 6.B.ii–iii), and cells further away from these growth factors were more stationary (red tracks). Both trends were not present in donor 1’s StemSpan microchip, giving credence that SCF and TPO enhanced HSPC proliferation and migration within 200 µm of inclusion, though it was found that migration preferentially occurred in the direction of growth factor sources.

While this system demonstrates the ability to evaluate HSPC proliferative response to a particular distribution of four growth factors, space within the chip occupied by these growth factors in its current configurations appears too frequently below their effective concertation throughout culture. Particularly, the ANG-2 and IGF-II, which exhibit maximal diffusion distance above their effective concentrations only 20% and 7% of the device distance, respectively, consequently have minimal overlap above effective concentration with each other, TPO and SCF environments. In future studies, the device dimensions could correspond precisely to the minimum diffusion distance of the least distributed factor. In this instance, a rectangle of 0.54 mm (7% of 7.64 mm) and 4.58 mm (60% of 7.64 mm) with SCF/TPO on the short axis and ANG-2/IGF-II on the long axis could have more consistently produced concentration ranges distributed above the effective concentration of all four molecules. This limited accessibility of cells to growth factors in the devices current configuration is potentially impacting the ability of the culture environment to maintain CD34^+^ HSPCs and prevent their differentiation and/or apoptosis, therefore resulting in impaired ability to study HSPC expansion.

Additionally, the scope of the presented study explores only a single growth factor in each of the four channels, which limits the possible concentration combinations of factors distributed spatially across the device. One possible solution could be to add three of the four factors to each channel, omitting a different factor in each channel. This would allow cells to be more consistently exposed to three of the four factors at all times, whilst also allowing the ability to study the influence of omitting a particular factor in combination with the remaining three factors. To even further increase the possible combinations of spatially distributed factors, it would be necessary to invert the media channel formulations in adjacent channels. As, the current device design is limited to four boundary conditions, to further improve the complexity of the system to study additional media factors and media formulations, projected polygons with additional sides could also be studied (e.g., pentagon, hexagon, decagon etc.).

The ability to phenotypically characterise this culture system is currently a limitation. While the system is designed to limit the diffusion of growth factor proteins, it coincidentally, in its current format, also limits the diffusion of antibodies necessary for immunocytochemical characterisation. A solution could be inspired by the reversible PDMS-glass bonding strategy adopted by Georgescu et al which could similarly be used allow directed access to the hydrogel culture and reduce the distance required for diffusion of antibodies and dyes^[17]^. Recovery of cells from the culture systems via enzymatic digestion of hydrogel and characterisation approaches such as quantitative polymerise chain reaction and flow cytometry could also benefit from this strategy to further characterise hydrogel-expanded cells and also give insight into any impact of the cell recovery process for future cell therapy application.

We validated the capability of this system to evaluate the proliferative response of CD34^+^ cells to induced gradients of SCF, TPO, ANG-2 and IGF-II. While it was hypothesized that the factors chosen could be supportive of fibrin hydrogel-based HSPC maintenance and proliferation, using the chosen geometry of 7.64 × 7.64 × 0.05, this environment appeared to mostly induce CD34^+^ erythroid differentiation. It should be highlighted that this lack of hypothesised cell maintenance is not just accredited to the chosen factors, but also their spatial distribution across the system and the choice of hydrogel. Interestingly, (non-gradient) Stem Line II cultures also demonstrated a large proportion of CD235a-positive cells, potentially alluding to the influence of the fibrin hydrogel on the cell phenotype. Nonetheless, this system demonstrates great potential in evaluating and identifying the type and concertation of culture media growth factors for hydrogel-based culture of CD34^+^ cells.

## 3. Conclusion

We demonstrate a platform technology capable of tracking hydrogel-based cell proliferation and migration in the presence of experimental growth factor gradients. The response of human UCB-derived CD34^+^ cells to quantified gradients of human recombinant growth factors TPO, SCF, ANG-2 and IGF-II demonstrated regionally asymmetrical expansion and factor preference, though further investigation is required. This technology exhibits potential in evaluating culture growth factors towards identifying media formulations for expanding or differentiating cells for applications in therapeutic cell manufacturing.

## 4. Experimental Section

### 4.1. Microfluidic device design, fabrication and preparation

Microfluidic devices were prepared as previously described in detail by Murphy et al (2025).^[29]^ Briefly, silicon/SU-8 master wafers were designed in AutoCAD (Autodesk, Inc.) 3D design software and fabricated by the Australian National Fabrication Facility Queensland Node to a feature height of 50 µm (figure 1.A.i). Silicon master wafers were rendered hydrophilic via overnight surface treatment with the vapour of hexamethyldisilizane.

Using SYLGARD 184 Silicone Elastomer Kit (Dow), 10-parts monomer was added to 1-part co-monomer, simultaneously mixed and degassed, and poured onto the master wafer to a height of approximately 5 mm above the wafer. The mixture was cured at 80 °C for a minimum of 1 hour before being carefully removed using a scalpel. Media reservoirs and cell injection ports were created using 1.25- and 4-mm diameter biopsy punches, respectively. The monolith elastomer and a rectangular glass coverslip were then treated with oxygen plasma at 35 W for 40 seconds before being irreversibly bonded via direct contact.

Microfluidic devices were wet autoclave sterilised at 121 °C before being washed of any residue via complete immersion in 70% (w/v) ethanol for a minimum of 1 hour. Ethanol was then completely removed from the devices via vacuum aspiration before the devices were washed in triplicate with sterile distilled water (15230162, Gibco^TM^). Devices were finally dried overnight in a humidified incubator at 37 °C in 5% (v/v) CO_2_ in air to remove any remaining liquid residue from channels. Devices were placed into 4-well rectangular culture dishes (167063, Thermo Fisher Scientific) with the middle two wells occupied by microfluidic devices (two devices per well) and the outer two wells filled with sterile distilled water or Dulbecco’s Phosphate Buffered Saline (DPBS, Gibco™) to prevent evaporation during culture.

### 4.2. Growth factor distribution modelling

Idealized 2D growth factor distributions were simulated within the hydrogel compartment of the microfluidic device. Following Altan-Bonnet & Mukherjee, growth factors were assumed to degrade throughout culture via half-life decay and transport from microfluidic channels into the hydrogel culture compartment via diffusion.^[30]^ This 2D approach assumes negligible spatial gradients exist in the *z-*dimension of the hydrogel compartment, which is reasonable due to the hydrogel chamber’s thin height (50 µm) so nearly all cells were cultured along one monolayer. A similar 2D simulation approach has been applied to model other thin tissue culture constructs^[29,31]^.

This model assumes growth factor media supplementation and half-life decay are significantly greater than other kinetics, such as growth factor secretion and cell or matrix sequestering, as found in prior work modelling single-cell cytokine release and binding^[25,32]^. Therefore, 2D growth factor transport and decay over time within the gradient microfluidic device were simply modelled as:

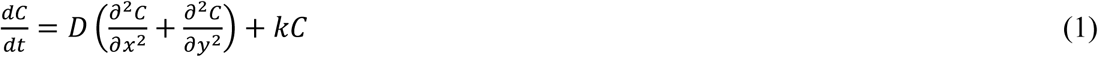

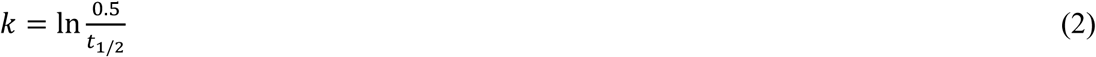

where, growth factor concentration *C* is modelled as a partial differential equation of dimensions *x*, *y* and *t* (equation 1). Here, *D* represents a growth factor’s diffusivity constant (μm^2^ s^−1^) and *k* is half-life decay rate (equation 2) with *t*_1/2_ being the time for the growth factor to decay to half its original concentration. This simulation applies the same half-life decay to the microchannel boundary concentration conditions, as media is only replenished every 2–3 days. The device’s 3D hydrogel cell culture compartment is 7.64 mm wide and 7.64 mm long, where four media channels interface with the culture chamber at its top-left, top-right, bottom-left, and bottom-right corners for 7.64 mm of length, each (figure 1.A.i). Table 1 lists major growth factors’ initial conditions, diffusivity, half-life, half maximal effective concentration (EC50) from these experiments. These parameters were derived from 2D culture conditions and growth factor transport, decay and effect would likely change in hydrogels.^[33]^ The distribution of each growth factor (ANG-2, IGF-II, SCF or TPO) was individually simulated across the microfluidic cell culture compartment (figure 1.A.ii, figure S4). The Python simulation code is provided in the supplementary information for an exemplary pair of counter-current growth.

**Table 1.** Growth factor distribution modelling parameters. FL: fetal liver, BM: bone marrow. ANG-2: angiopoietin 2, IGF-II: insulin-like growth factor II, SCF: stem cell factor, TPO: thrombopoietin.

| HSC niche | Growth Factor and References | Supplemented Concentration [ng mL<sup>-1</sup>] | Diffusivity [$\mu\text{m}^2 \text{s}^{-1}$] | Half-life [s] | Half-effective Concentration [EC50; ng mL<sup>-1</sup>] |
| --- | --- | --- | --- | --- | --- |
| FL | ANG-2<sup>[34,35]</sup> | 100 | 13.3 | 10,800 | 0.5–4 |
| FL | IGF-II<sup>[36,37]</sup> | 100 | 30 | 1,260 | 1.5–6 |
| FL/BM | SCF<sup>[38]</sup> | 100 | 121 | 170,000 | 1–5 |
| FL/BM | TPO<sup>[39–41]</sup> | 100 | 98 | 108,000 | 0.3–3 |

### 4.3. Mononuclear cell fraction separation and CD34^+^ cell isolation

This project was supported by The University of Queensland’s (UQ) Human Research Ethics Committee through approval 2022/HE001310, as well as UQ’s Institutional Biosafety Committee through approval IBC/568B/ChemEng/2022.

#### 4.3.1. Density centrifugation

Human UCB-derived buffy coat was obtained from the Queensland Cord Blood Bank At The Mater (South Brisbane, Queensland, Australia) following validated donor education and informed consent, *ex utero* cord blood collection and volume reduction via centrifugation and using the MacoPress SMART (MacoPharma, France) procedures. Biological starting material was provided for research with appropriate consent under materials transfer agreement CBB-QF-LP-027.2. Buffy coats were transported at room temperature to the UQ for CD34^+^ cell isolation. A total of four UCB donations were used in this study. After gently mixing by hand, blood bags were drained and approximately 25 mL of buffy coat was equally transferred to two 50 mL centrifuge tubes (12.5 mL per tube). Blood was then diluted 1:1 (v/v) with Hanks Balanced Saline Solution (HBSS, H6648, Sigma-Aldrich) containing 0.6% (v/v) citrate-dextrose solution (ACD, C3821, Sigma-Aldrich). The diluted blood was then carefully (as to not mix the two) laid over 20 mL of room temperature Histopaque®-1077 Hybri-Max™ (H8889, Sigma-Aldrich) and centrifuged at 400×g for 30 minutes at room temperature (acceleration: 6, deceleration: 5). The top clear-yellow plasma layer was then aspirated being careful not to disturb the middle mononuclear cell (MNC) interphase. The MNC interphase was then collected into a 50 mL centrifuge tube to which 45 mL of HBSS was added, the cap secured and mixed via gentle inversion. This cell suspension was then centrifuged at 250×g for 10 minutes at room temperature (acceleration: 9, deceleration: 4), supernatant discarded, and cells again resuspended 45 mL HBSS. This washing procedure was repeated up to 6-times or until the supernatant was no longer visibly cloudy. The cell pellet was then resuspended in 5 mL of 37 °C red blood cell lysis buffer (RBCLB, 150 mM NH_4_Cl, 1 mM K_2_EDTA.2H_2_O, 10 mM KHCO_3_; pH 7.5), mixed gently via manual inversion and incubated at room temperature for 5 minutes. While lysing, a 20 µL sample was taken, diluted 1:100 in RBCLB, of which, 20 µL was mixed with 20 µL 0.04% trypan blue in HBSS and counted either manually using a hemacytometer or automatically using a Countess II automated cell counter. After incubation, the cell mixture was then diluted to 50 mL with 0.6% ACD in HBSS and centrifuged at 200×g for 5 minutes at room temperature (acceleration: 9, deceleration: 4) to remove platelet contamination. The supernatant was then discarded, and the cell pellet resuspended in 20 mL HBSS.

#### 4.3.2. Immunomagnetic separation

The cell suspension from 4.3.1. was centrifuged at 200×g for 5 minutes at room temperature (acceleration: 9, deceleration: 4), supernatant aspirated and cell pellet resuspended in EasySep™ Buffer (20144, STEMCELL Technologies) at a concentration of 5 × 10^8^ cells mL^-1^ and transferred to a 14 mL polystyrene centrifuge tube (38008). The EasySep™ Human CD34 Positive Selection Cocktail (17856C, STEMCELL Technologies) was then added at 100 µL mL^-1^ of sample, mixed and incubated at room temperature for 10 minutes. The EasySep™ Dextran RapidSpheres™ (50100, STEMCELL Technologies) were vortex mixed for 30 seconds and added to the sample at 75 µL mL^-1^ of sample, mixed and incubated at room temperature for 5 minutes. The sample was then made up to 3 mL with EasySep™ Buffer, mixed gently and the tube (without lid) placed in the “The Big Easy” EasySep™ Magnet (18001, STEMCELL Technologies) for 3 minutes at room temperature. The supernatant was then discarded by pouring the tube (remaining in the magnet) into a 50 mL centrifuge tube. This process was repeated up to 4 more times or until supernatant became visibly clear. The tube walls were then washed in 3 mL of EasySep™ Buffer to collect the CD34^+^ cells. Cells were then counted as described in 4.3.1, then transferred to a 15 mL centrifuge tube and centrifuged at 200×g for 5 minutes at room temperature (acceleration: 9, deceleration: 9). Supernatant was discarded and cells resuspended in human AB serum (H4522, Sigma-Aldrich) with 10% v/v dimethylsulfoxide (D2438, Sigma-Aldrich) and cryopreserved in liquid nitrogen vapour.

### 4.4. CD34^+^ cell routine suspension expansion

CD34^+^ cells were thawed at 37 °C for approximately 5 minutes and transferred to 9 mL of human AB serum. The cell suspension was then centrifuged at 200×g for 5 minutes, supernatant aspirated and cell pellet gently resuspended in 0.5 mL of StemSpan™ SFEM II supplemented with 1X StemSpan™ CD34^+^ Expansion Supplement (10X; #02691, STEMCELL Technologies), herein referred to as ‘complete CD34^+^ cell expansion media’. To ensure cell viability, 0.4% w/v trypan blue stock solution was diluted to 0.04% in HBSS for cell counting. 10 µL of cell suspension was mixed with 10 µL of 0.04% w/v trypan blue working solution and a viable cell count was conducted manually using a hemocytometer or automatically using a Countess II automated cell counter. The cell suspension was then diluted to a final concentration of 10^4^ viable cells per mL in complete CD34^+^ expansion media in an appropriately sized culture vessel in a humidified incubator at 37 °C in 5% (v/v) CO_2_ in air. Cultures were left to expand for 4 days without media exchange or dilution before being harvested for microfluidic culture experiments.

### 4.5. Hydrogel-based microfluidic gradient cell culture

#### 4.5.1. Reagent preparation

##### Hydrogel components

Fibrinogen type 1-S from bovine plasma (F8630, Sigma-Aldrich) stock solution was prepared as per manufacturer instructions at 10 mg mL^−1^ in HBSS before being sterilized using a 0.22 μm syringe filter. Thrombin from bovine plasma (T7513, Sigma-Aldrich) stock solution was prepared at 100 U mL^−1^ in 0.1% (w/v) bovine serum albumin solution and similarly sterilized. Aprotinin (A1153, Sigma-Aldrich) was dissolved in DPBS to give a stock solution of 15 U mL^−1^ and sterile filtered.

##### Growth factor reconstitution

Recombinant human angiopoietin-2 protein (ANG-2; 623-AN, R&D Systems), recombinant human insulin-like growth factor II protein (IGF-II; 292-G2, R&D Systems) and recombinant human stem cell factor (SCF; S7091, Sigma-Aldrich) were all reconstituted at 100 µg/mL in sterile DPBS with 0.1% w/v bovine serum albumin. Recombinant human thrombopoietin (TPO; T1568, Sigma-Aldrich) was first reconstituted at 500 µg/mL in 10 mM sodium citrate pH 3 then diluted to 20 µg/mL in sterile DPBS with 0.1% w/v bovine serum albumin.

##### Gradient media preparation

For gradient cultures, Stem Line II Hematopoietic Stem Cell Expansion Medium without antibiotics or cytokines (S0192, Sigma-Aldrich) was supplemented with 100 ng mL^-1^ of either ANG-2, IGF-II, SCF or TPO. For homogenous control cultures, a mixture of the four factors at 100 ng/mL each was prepared.

#### 4.5.2. Microfluidic culture

Fibrinogen stock solution was diluted to 6 mg mL^−1^ in HBSS. Cultured CD34^+^ cell suspensions were centrifuged at 200×g for 5 minutes and resuspended at 20 × 10^6^ cells mL^−1^ using a solution of 2 U mL^−1^ thrombin and 0.3 U mL^−1^ aprotinin in complete CD34^+^ cell expansion media. 10 μL of cell suspension was quickly mixed with 10 μL of 2 mg mL^−1^ fibrinogen, then 10 μL of this mixture (1 mg mL^−1^ fibrinogen, 1 U mL^−1^ thrombin, 0.15 U mL^−1^ aprotinin) was pipetted directly into the cell seeding port of the microfluidic device. The cell-seeded device (10 × 10^6^ cells mL^−1^) was then placed in a humidified incubator for 15 minutes at 37 °C in 5% (v/v) CO_2_ in air to facilitate fibrin hydrogel formation. Media channels were then loaded with media (single growth factor media, 4-mixed growth factor media or CD34^+^ cell expansion media) and reservoirs filled with ≈75 μL similarly matched media with 0.15 U mL^-1^ aprotinin. Devices were finally placed in a humidified incubator at 37 °C in 5% (v/v) CO_2_ in air with complete media after approximately 1-day and 3-days post seeding before terminating at approximately 5-days.

### 4.6. Time lapse imaging and analysis

#### 4.6.1. Image acquisition

Time-lapse microscopy of microfluidic cultures was conducted using a Etaluma LS720 in-incubator fluorescence microscope with automated stage control (figure 1.B) running Lumaview 720/600-Series software. Images were acquired with minimum LED intensity to minimize phototoxicity and minimum photomultiplier gain to minimize image signal-to-noise. A 15 × 15 image grid with 10% image overlap was acquired across the complete 7640 × 7640 µm culture compartment of the microfluidic device using a 20× Olympus LUCPlan FLN NA 0.45 objective. Multipoint time-lapse protocols were conducted simultaneously across two or three microfluidic devices and run constantly (approximately 1 image per 1.33 seconds) over the course of up to 5-days culture. After media exchanges on days 2 and 3, imaging positions were reset due to small changes in the *xyz*-positions of the device.

#### 4.6.2. Image analysis

Full microchip tile scan images were stitched together as 15 × 15 grids with a 10% overlap by using an automated ImageJ FIJI macroscript. For a typical microchip culture span of 120-time steps between media exchanges, each with 450 (2-chips) to 675 (3-chips) brightfield images; this would require 1–2 days or 4–5 days for each microchip of a 5-day (two media exchange) culture and result in tiled images of roughly 1–2 gigabytes in size.

Tiled 15 × 15 imaged were then manually aligned and cropped to the cell culture chamber inside the trapezoid hydrogel pillars (7640 × 7640 µm size). Then, these images were analysed by the cellpose inbuilt ‘cyto’ cell segmentation model for greyscale brightfield images, provided a mean diameter of 31.9 pixels. Cellpose’s single-cell segmentation results were output to a table with cell centroids (X, Y), equivalent diameter, area, perimeter, and orientation, all in units of pixels. We then estimated circularity and local density through the following metrics:

- props[“Circularity”] = 4*math.pi*(props[“Area”]/props[“perimeter_crofton”]**2)

- props[“Local Density”] = gaussian_kde([props[’Centroid-0’],props[’Centroid-1’]])([props[’Centroid-0’],props[’Centroid-1’]])

While circularity had units between 0 and 1, where 1 represents a perfect circle; the units of local density, calculated as a kernel density estimation, sum to 1 across the entire image space. Splitting the microchip culture space into 1,000 × 1,000 regions, the local density was typically in the range of 10^-6^.

As discussed above, cell tracking was prototyped by asking, for each of the 20,000–150,000 of cell centroids detected within each of the 300–500 imaged timesteps, whether the next timestep presents a cell centroid inside of a 150 µm Euclidean distance radius of that cell in the prior timestep. And if multiple future cell centroids were detected to be close to that past cell centroid, the closest future cell was selected as the past cell’s next tracked position. If no future cell centroids were detected in that threshold radius, the cell was considered to have died. If a new cell was detected and not related to a past cell, it was considered a newly proliferated cell.^[42]^

### 4.7. Immunofluorescence staining and imaging

Cultures were fixed at room temperature with 3.7% (w/v) paraformaldehyde (15 710, ProSciTech) for up to 2 hours then washed with DPBS three times for 15 minutes each. Fixed cultures were then permeabilized with 0.1% (w/v) Triton X-100 solution in DPBS for up to 2 hours followed by three 15-minutes washes with DPBS, then incubated in 10% (v/v) normal goat serum (NGS, 50062Z, Thermo Fisher Scientific) in DPBS blocking solution at 4 °C for 48 hours on a rocker platform (rotating the device 90° in the *xy*-plane after 24 hours). Cultures were then incubated in anti-CD90 IgG (MA5-32559), anti-CD31 IgG2a (MA3100) and respective isotype control primary antibodies (31 235, 14-4724-82 and 14-4724-82), diluted in 10% (v/v) NGS, at 4 °C for 48 hours on a rocker platform (rotating the device 90° at 24 hours), followed by three 15-minute washes with DPBS. Secondary antibodies anti-rabbit IgG AlexaFluor488 (A-11008), anti-rat IgG AlexaFluor555 (A-21434), anti-mouse IgG2a AlexaFluor555 (A-21137), anti-rabbit IgG AlexaFluor647 (A21245), anti-mouse IgG2a AlexaFluor647 (A-21241) and anti-mouse IgG1 AlexaFluor647 (A-21240) were diluted to 4 μg mL^−1^ in 10% (v/v) NGS followed by another three 15-minute washes with DPBS. 4′,6-diamidino-2-phenylindole, dihydrochloride (DAPI) stock solution (R37606, Thermo Fisher Scientific) diluted 2 drops mL^−1^ in DPBS was applied to cultures and incubated for 30 minutes at room temperature, followed by another three 5-minutes washes with DPBS. All primary and secondary antibodies were purchased from Thermo Fisher Scientific. Images were obtained using a Nikon Ti-2 inverted fluorescence microscope running NIS Elements software.

## Acknowledgements

The authors would like to thank Dr Rebecca E. Griffiths and Coleen Venida of the Australian Red Cross Lifeblood for sharing their HSC isolation protocol and providing training.

This work was supported by funding from the University of Queensland Philanthropic Fund, the National Heart Foundation of Australia (108132-372024_FLF), the Australian Research Council (DE220100757), and a Ramaciotti Health Investment Grant (2022HIG08).

This work used the Queensland node of the NCRIS-enabled Australian National Fabrication Facility (ANFF).

## Data Availability Statement

The data that support the findings of this study are available from the corresponding author upon reasonable request.

## Supporting Information

Supporting Information is available from the Wiley Online Library or from the author.

Received: ((will be filled in by the editorial staff))

Revised: ((will be filled in by the editorial staff))

Published online: ((will be filled in by the editorial staff))

## Supporting Information

**Figure S1.**
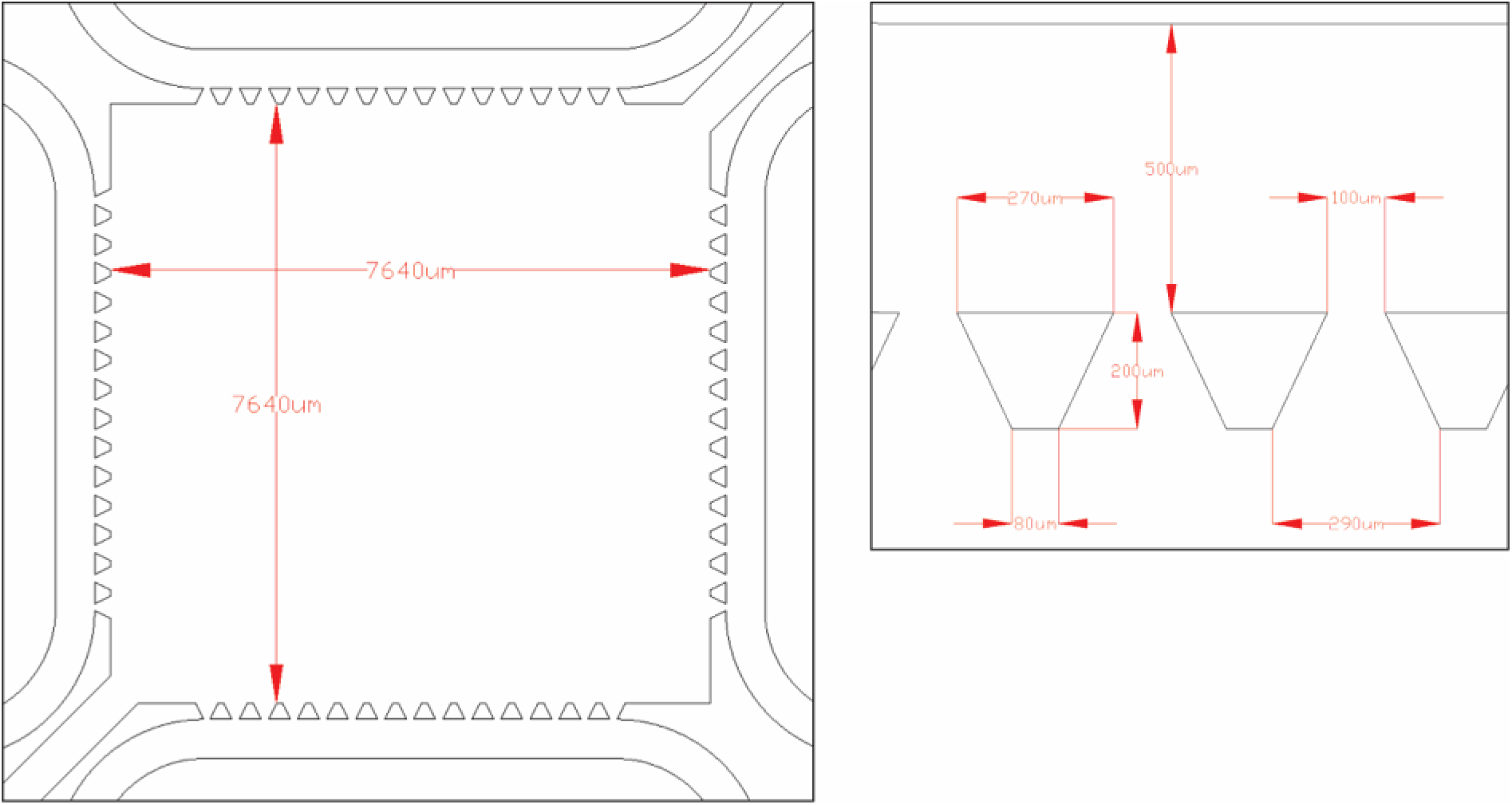
Microfluidic culture device dimensions.

**Figure S2.**
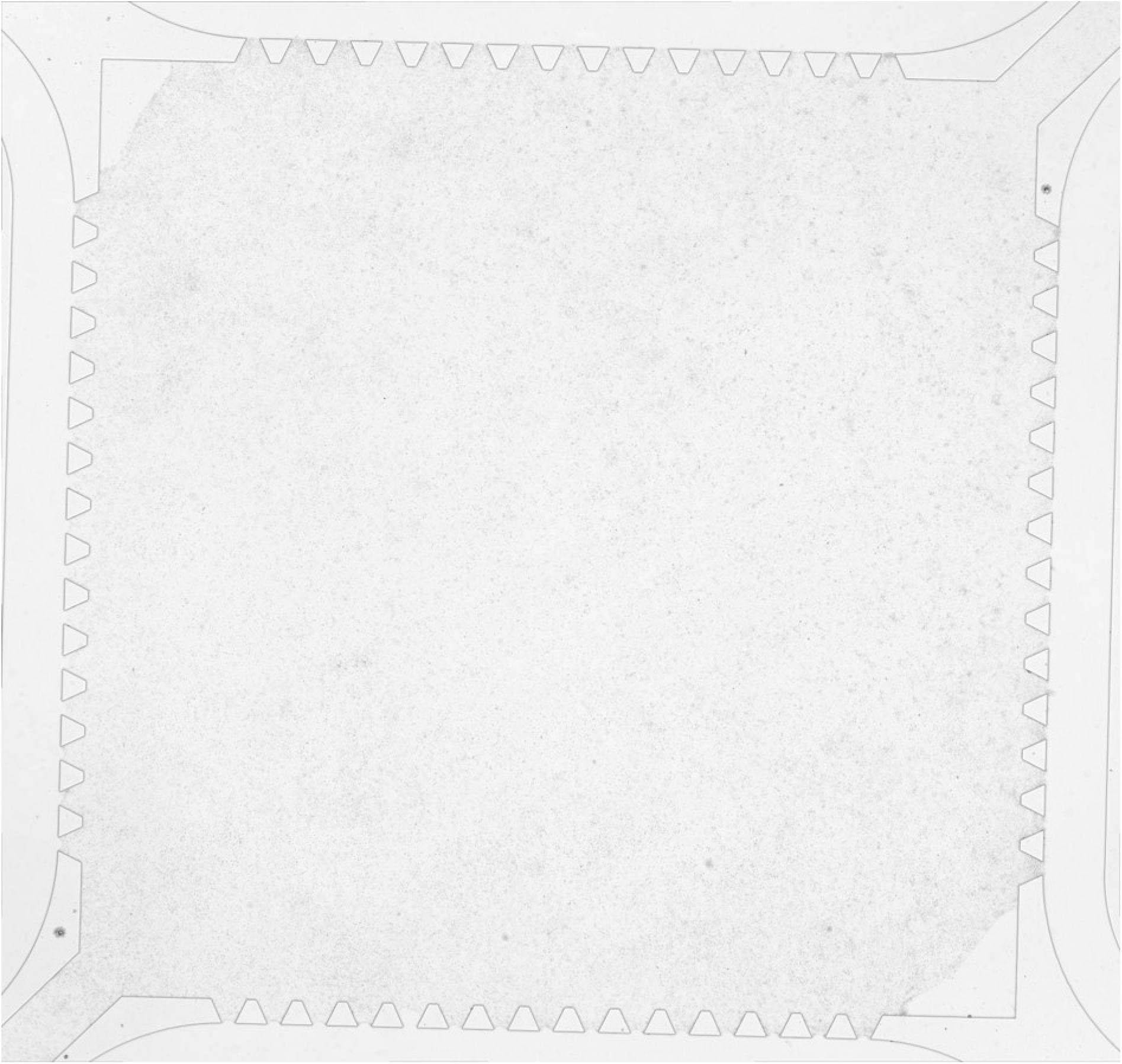
Microfluidic devices fill near completely within the culture compartment. Microfluidic chip loaded with 10 µL of hydrogel precursor cell suspension (1 mg mL^−1^ fibrinogen, 1 U mL^−1^ thrombin, 0.15 U mL^−1^ aprotinin; 10 × 10^6^ cells mL^−1^) extended to the edges of confining trapezoidal pillars but fails to completely fill the adjacent chip corners.

**Figure S3.**
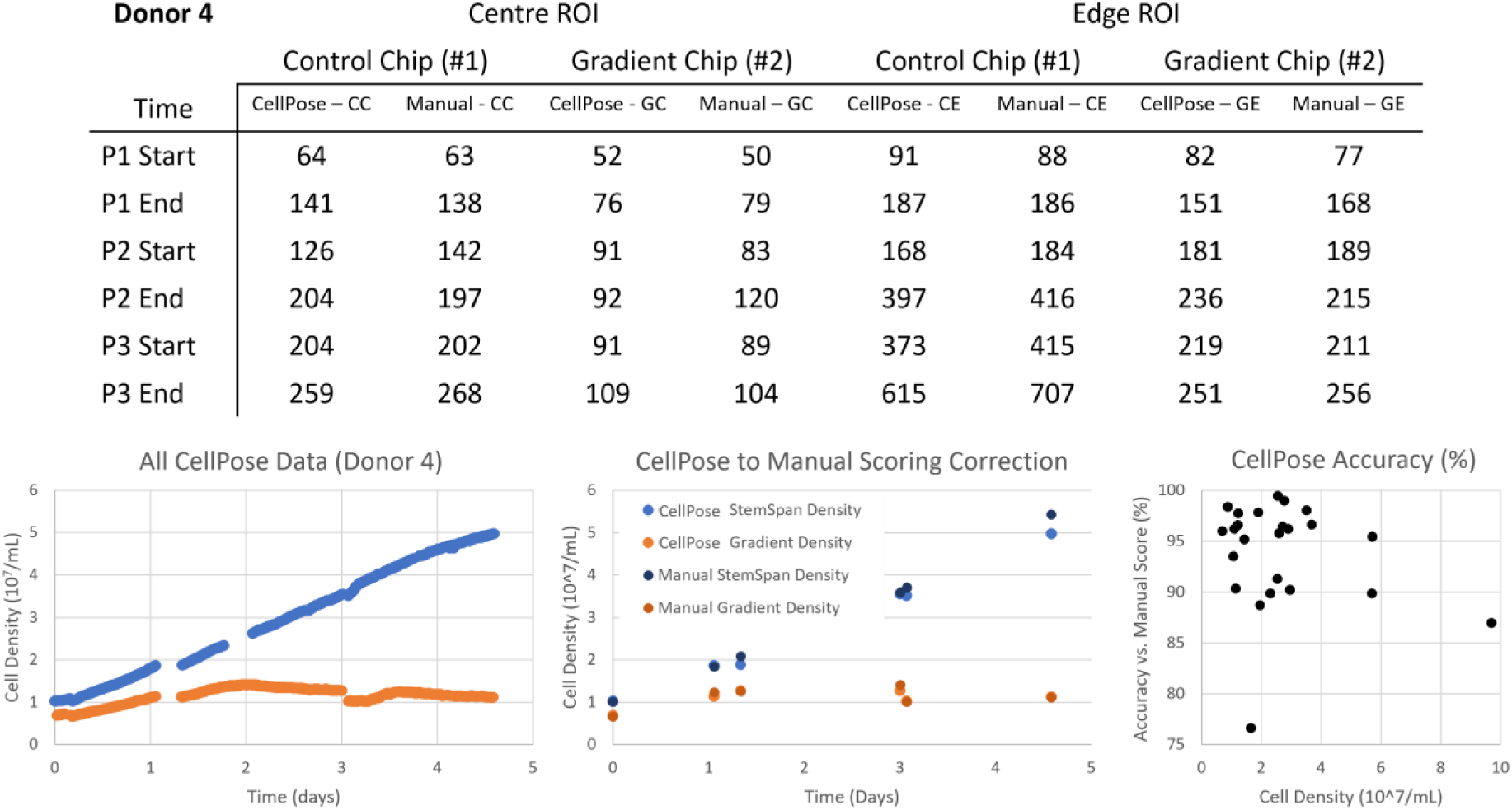
Cellpose single-cell detection accuracy. Regions of interest (ROIs) at 1/20^th^ the scale of the 15×15 full microchip tile scans were taken from the centre and right edge of the microchips at the beginning and the end of donor 4 three intervals between media exchanges (P1, P2, P3) in two types of microchips (StemSpan Control, Gradient). The 24 ROIs were manually counted for cells by an experienced bioengineer and compared to the cellpose detection from the full tile scan. Cellpose detections were found to be 93.86 ± 5.02% accurate over 4,647 cells in 24 manually scored images at varied densities. There did not seem to be a correlation in accuracy between culture time or culture density, however, it was evident that cultures with less viable cells (fragmented cell membranes), such as the centre of donor 4’s gradient chip, produced the lowest cellpose detection accuracy of 76.7%.

**Figure S4.**
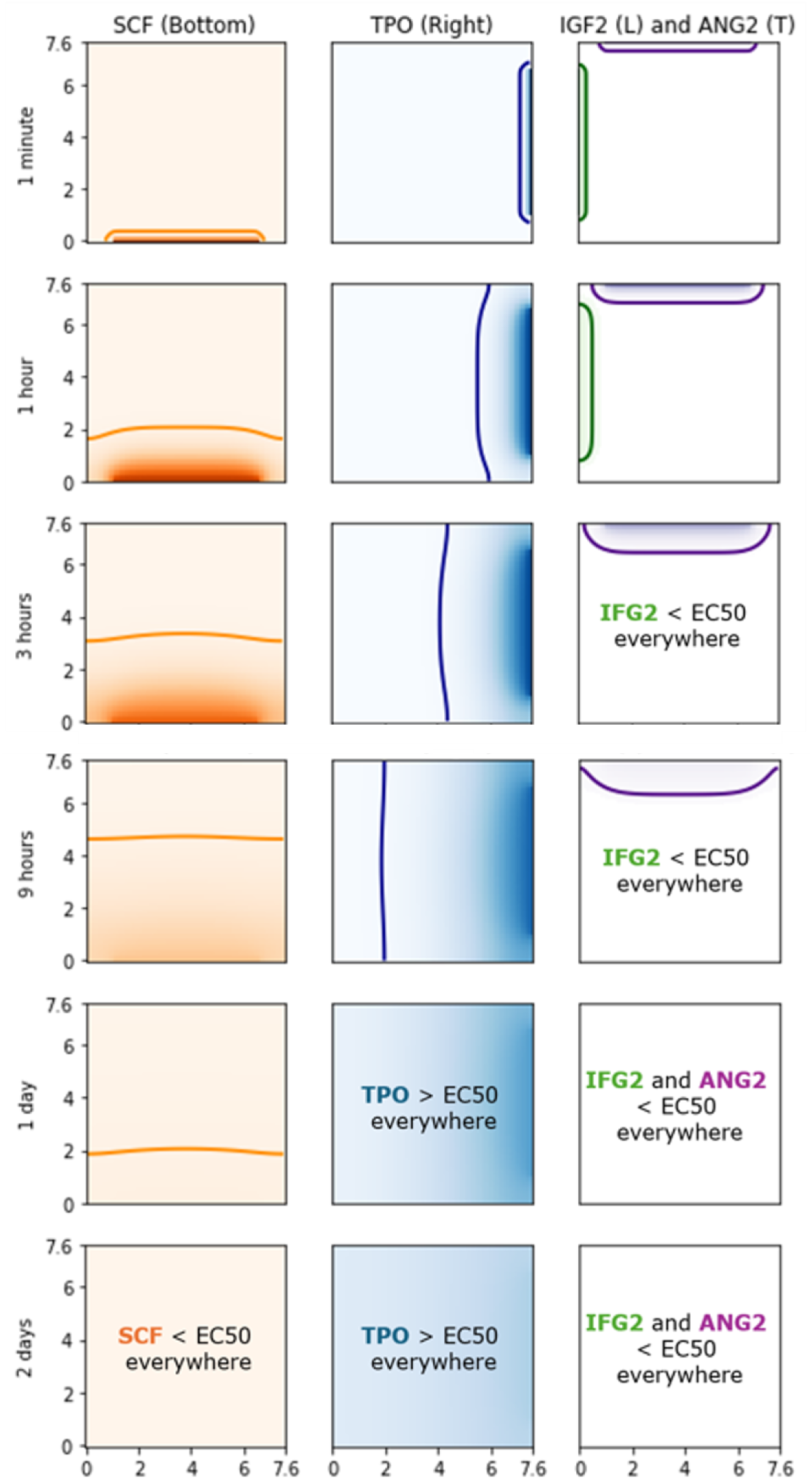
Diffusion-decay growth factor simulations in gradient microchips. Heatmaps represent relative concentration of SCF, TPO, IGF2, and ANG2 using diffusivity, half-life, and EC50 information from Table 1 and normalised to a maximum concentration of 100 ng mL^-1^. The overlaid contours represent where the diffusing growth factors reach their EC50 (ng/mL). IGF-II, ANG-2, and SCF do not exist above their EC50 value from hours 3, 9, and 48, respectively, while TPO is above its EC50 value throughout the entire chip from 24-hours post seeding.

**Figure S5.**
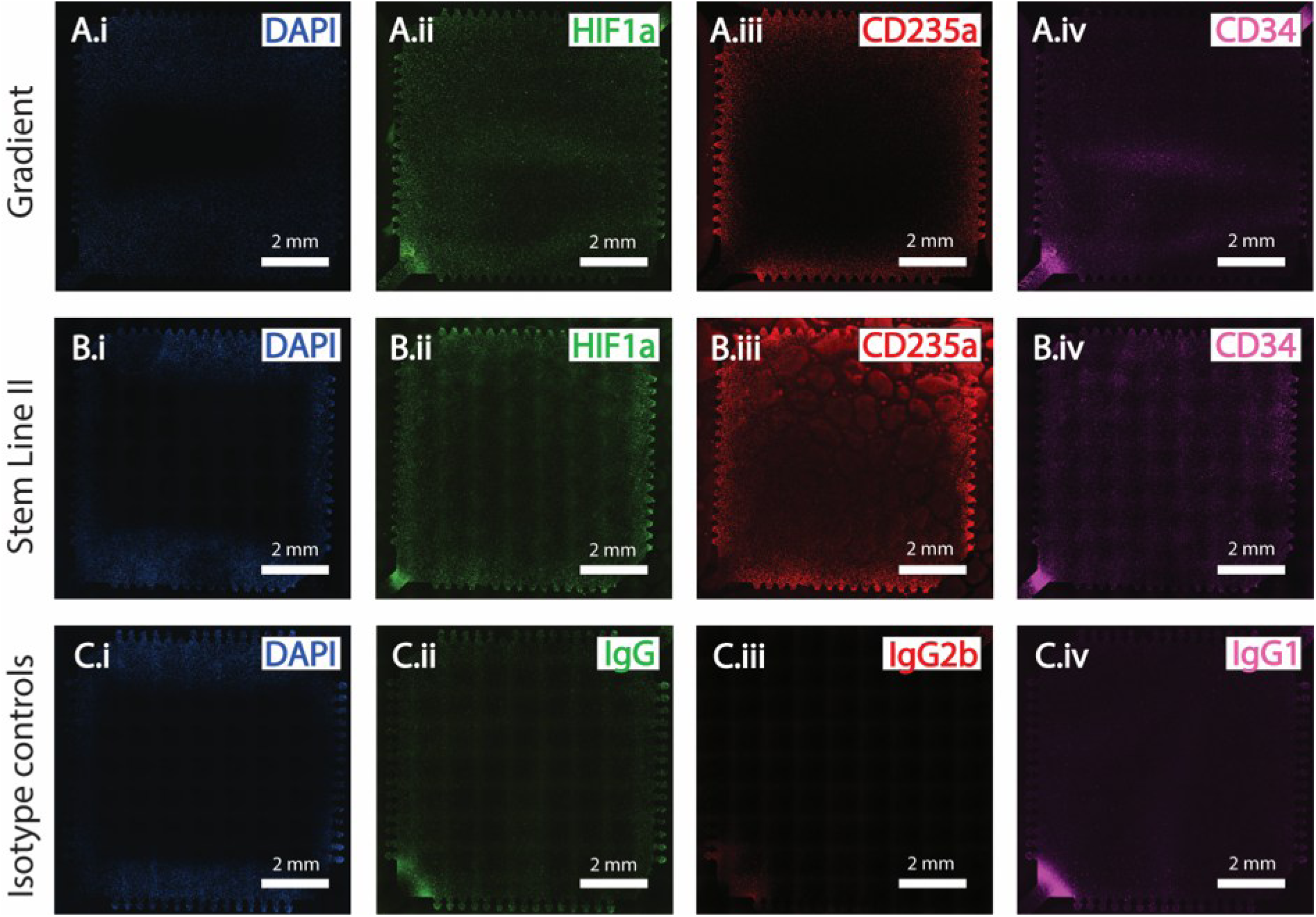
Fibrin hydrogel-based cells express markers of cellular stress, hematopoietic maintenance and hematopoietic lineage after 5-days culture. Fibrin hydrogel-based gradient (**A**) and Stem Line II (**B**) cultures were fixed at day-5 post seeding and nuclear counterstained with (*i*) DAPI and immunocytochemically stained for the expression of (*ii*) HIF1a, (*iii*) CD235a and (*iv*) CD34. **C.** Respective isotype control stains. Scale bars = 2 mm.

**Figure S6.**
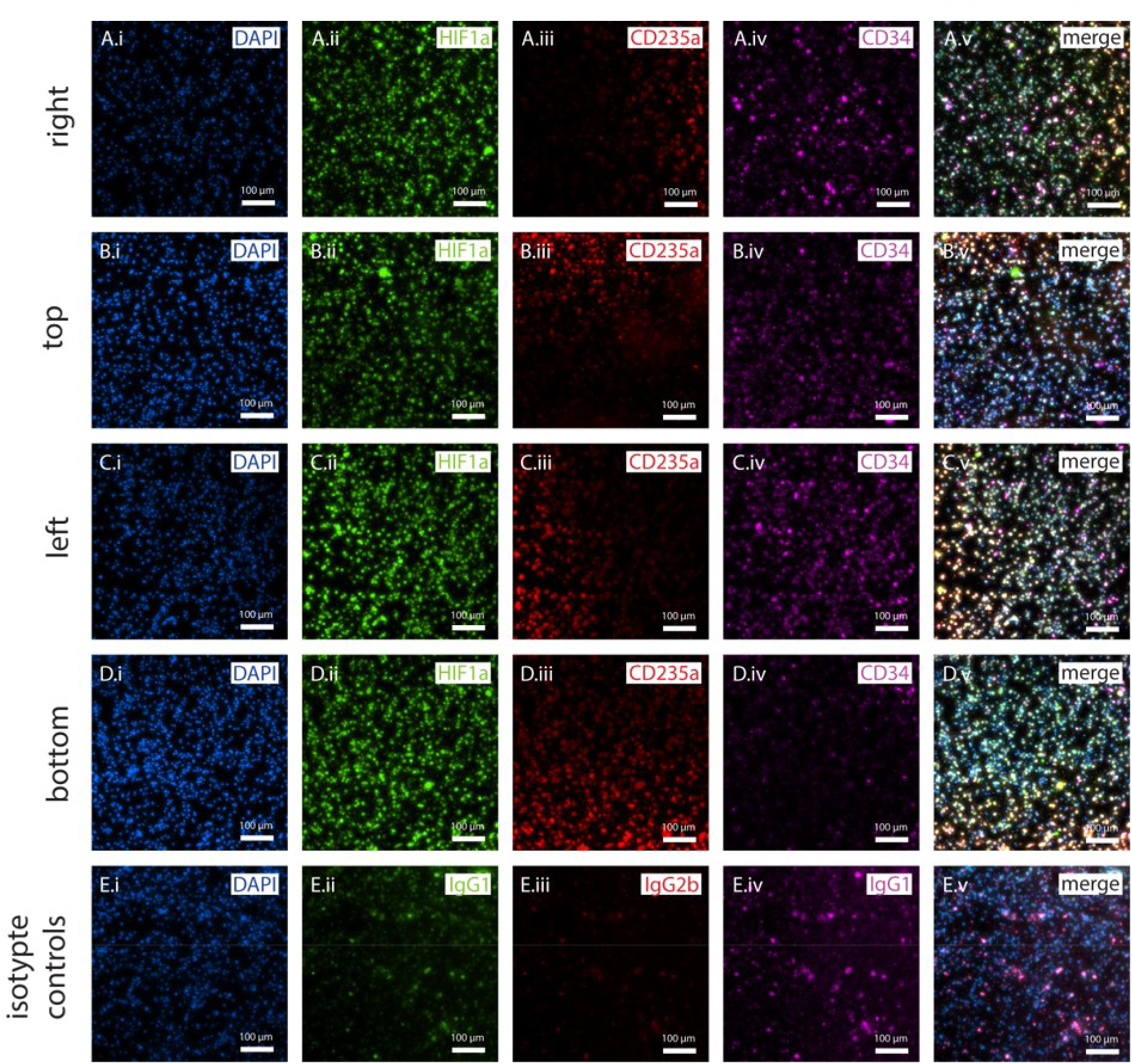
Stem Line II fibrin hydrogel-based cells express markers of cellular stress, hematopoietic maintenance and hematopoietic lineage after 5-days culture. Fibrin hydrogel-based Stem Line II cultures were fixed at day-5 post seeding and nuclear counterstained with (*i*) DAPI and immunocytochemically stained for the expression of (*ii*) HIF1a, (*iii*) CD235a and (*iv*) CD34. Images were capture at the edge of the hydrogel near the right (**A**), top (**B**), left (**C**) and bottom sides (**D**). **E.** Respective isotype control stains. Scale bars = 2 mm.

**Table S1.** Average normalised cell diameter changes over the course of fibrin hydrogel-based CD34^+^ microfluidic cell culture. Normalised cell diameter across the 5-days of culture (T_0_ ≈ 2 hours, T_f_ ≈ 5 days). f: final.

| Donor | Condition | $T_0$ | | $T_f$ | |
| --- | --- | --- | --- | --- | --- |
|  |  | *Avg* | *Std* | *Avg* | *Std* |
| Donor 1 | Stem Line II | 24.33 | 17.17 | 21.57 | 15.48 |
| Donor 1 | Gradient | 24.38 | 17.43 | 23.03 | 17.47 |
| Donor 2 | StemSpan | 22.69 | 13.85 | 17.61 | 13.24 |
| Donor 2 | Stem Line II | 23.30 | 14.66 | 17.14 | 12.84 |
| Donor 2 | Gradient | 23.16 | 14.27 | 17.77 | 13.14 |
| Donor 3 | StemSpan | 23.16 | 13.99 | 17.76 | 12.65 |
| Donor 3 | Stem Line II | 23.99 | 14.51 | 18.09 | 12.26 |
| Donor 3 | Gradient | 22.46 | 13.85 | 17.22 | 12.87 |
| Donor 4 | StemSpan | 22.32 | 12.84 | 19.56 | 13.35 |
| Donor 4 | Gradient | 22.05 | 12.56 | 18.15 | 13.53 |

**Video S1. Time-lapse imaging captures of fibrin hydrogel-based CD34^+^ cell culture.** CD34^+^ cells seeded at a density of 10 × 10^6^ cell mL^-1^ in 1 mg ml^-1^ bovine fibrin hydrogels within a culture compartment volume of 2.92 µL and cultured up to 5-days. Time lapse images of donor 4 StemSpan condition region of interest 60. Discontinuations in data represent culture media exchanges and resetting of the *xyz*-imaging coordinates. Images captured approximately every 8 minutes. Field of view is 670 × 670 µm. *[video file attached to submission]*

**Video S2. Time-lapse imaging and quantitative cell segmentation captures of fibrin hydrogel-based CD34^+^ cell culture density in response to Gradient conditions.** CD34^+^ cells seeded at a density of 10 × 10^6^ cell mL^-1^ in 1 mg ml^-1^ bovine fibrin hydrogels within a culture compartment volume of 2.92 μL and cultured up to 5-days. Cells were detected via the cellpose software and local cell density qualitatively displayed with dark blue representing the lowest local cell density in a particular culture image set and yellow representing highest local cell density, for donor 4. The left panel represents the entire culture compartment (7.64 × 7.64 mm), the middle panel a single region of interest from the centre of the culture compartment (670 × 670 μm) and the right panel a single region of interest from the bottom right corner of the culture compartment (670 × 670 μm). Discontinuations in video data represent culture media exchanges and resetting of the *xyz*-imaging coordinates. Count = cell number. Time represents days:hours:minutes. *[video file attached to submission]*

**Video S3. Time-lapse imaging and quantitative cell segmentation captures of fibrin hydrogel-based CD34^+^ cell culture density in response to StemSpan conditions.** CD34^+^ cells seeded at a density of 10 × 10^6^ cell mL^-1^ in 1 mg ml^-1^ bovine fibrin hydrogels within a culture compartment volume of 2.92 μL and cultured up to 5-days. Cells were detected via the cellpose software and local cell density qualitatively displayed with dark blue representing the lowest local cell density in a particular culture image set and yellow representing highest local cell density, for donor 4. The left panel represents the entire culture compartment (7.64 × 7.64 mm), the middle panel a single region of interest from the centre of the culture compartment (670 × 670 μm) and the right panel a single region of interest from the bottom right corner of the culture compartment (670 × 670 μm). Discontinuations in video data represent culture media exchanges and resetting of the *xyz*-imaging coordinates. Count = cell number. Time represents days:hours:minutes. *[video file attached to submission]*

